# Scleral appearance is not a correlate of domestication in mammals

**DOI:** 10.1101/2023.02.27.530332

**Authors:** Kai R. Caspar, Lisa Hüttner, Sabine Begall

## Abstract

Numerous hypotheses try to explain the unusual appearance of the human eye with its bright sclera and transparent conjunctiva and how it could have evolved from a dark-eyed phenotype, as is present in many non-human primates. Recently, it has been argued that pigmentation defects induced by self-domestication may have led to bright-eyed ocular phenotypes in humans and some other primate lineages, such as marmosets. However, it has never been systematically studied whether actual domesticated mammals consistently deviate from wild mammals in regard to their conjunctival pigmentation and if this trait might therefore be part of a domestication syndrome. Here, we test this idea by drawing phylogenetically informed comparisons from a photographic dataset spanning 13 domesticated mammal species and their closest living wild relatives. We did not recover significant differences in scleral appearance or irido-scleral contrast between domesticated and wild forms, suggesting that conjunctival depigmentation, unlike cutaneous pigmentation disorders, is not a general correlate of domestication. Regardless of their domestication status, macroscopically depigmented conjunctivae were observed in carnivorans and lagomorphs, whereas ungulates generally displayed darker eyes. Based on our dataset, we also present preliminary evidence for a general increase of conjunctival pigmentation with eye size in mammals. Our findings suggest that conjunctival depigmentation in humans is not a byproduct of self-domestication, even if we assume that our species has undergone such a process in its recent evolutionary history.

## Introduction

The external appearance of mammalian eyes is highly variable. In addition to the shape of the pupil [1] and the coloration of the iris [2], it is importantly determined by the pigmentation of the bulbar conjunctival epithelium [3], which adheres to the externally visible portions of the sclera and is the focus of this study. The opaque scleral tissue has a white to greyish complexion, while the appearance of the conjunctiva can range from translucent to black dependent on the degree of melanin pigmentation [3]. Together, these structures create a phenotypic impression, which we will refer to here as “scleral appearance”. Traditionally, the important role of the conjunctiva in determining ocular complexion has been downplayed, with many authors simply referring to “the sclera” when discussing scleral appearance [4–8].

In humans, the sclera is bright (although it harbors substantial populations of melanocytes – [9]) and the overlying conjunctiva is transparent, creating the conspicuous white of the eye. This feature has attracted considerable research attention because it is commonly assumed to be rare among mammalian species, particularly primates [4–6, 10, 11]. Its phylogenetic origins and potential adaptive value, including communicative functions, are discussed extensively in the contemporary literature [6–8, 12–14]. In other hominid primates, the conjunctiva is also often partially depigmented and scleral appearance may be bright, but unlike in humans, a great intraspecific variation in this trait is typically evident [6, 14]. However, interindividually uniform white scleral appearance, resembling the human phenotype, can nevertheless be found in some monkey species, such as marmosets [7, 12, 15].

It has been suggested that the lack of macroscopically visible pigment in the conjunctivae of humans and some other primates is a byproduct of so-called self-domestication [7, 11, 16]. This concept assumes that selection against aggression in wild animal species, can give rise to a suite of traits that are otherwise characteristic of domesticated lineages [17]. These may include smaller brains and more delicate jaws, as well as pigmentation defects, resulting in the pied coat patterns of many domestic mammals [18]. The co-emergence of these different characteristics is commonly denoted as the domestication syndrome, which is assumed to affect both actual domesticated groups and wild self-domesticated lineages that have been subjected to similar selection pressures during their evolutionary history [16, 19]. The domestication syndrome is hypothesized to derive from pleiotropy: selection for tameness and reduced reactive aggression affects the adrenal glands, which secrete stress hormones into the bloodstream. In domesticated lineages, the adrenal glands are typically reduced in size, and stress hormone levels are markedly lower than in wild mammal species [17, 20]. Embryologically, the adrenal medulla derives from neural crest cells, a distinct population of stem cells that migrate through the embryo and develop into numerous tissues, including parts of the facial skeleton [21], various ocular structures [22], and the melanocytes of the integument [23]. Hence, via altering the migration of neural crest cells and their differentiation in the adrenal medulla, morphological changes in other neural crest-derived structures may follow as a byproduct of selection for tameness. Although there is compelling genetic evidence for pronounced alterations of the neural crest being correlated with domestication across mammalian lineages [24], no consensus has been reached on exactly which traits are comprised by the domestication syndrome. A recent review found that, apart from tameness and docility, skin depigmentation is the only trait that is universally expressed in domesticated mammals, calling the scope of the syndrome into question [18].

Whether conjunctival depigmentation is also affected by domestication and thus may represent a facet of the domestication syndrome has not yet been comparatively assessed. The ocular melanocytes of the uvea and sclera derive from a distinct lineage of neural crest cells that may be subject to different genetic regulations than melanocyte precursors that migrate into the skin [25]. Unfortunately, the precise embryological origins and maturation patterns of conjunctival melanocytes have apparently never been systematically studied (compare [26]). Just like integumental melanocytes and unlike those of the uvea and sclera, the melanocytes of the conjunctiva transfer pigment granules to adjacent cells [27]. Whether this is the consequence of close developmental ties remains to be clarified.

In any case, it is crucial to assess ocular pigmentation in domesticated and closely related wild mammals before one can make informed conclusions about whether conjunctival depigmentation in species such as humans, bonobos, and marmosets is indeed a correlate of self-domestication. It is also important to point out that the evolution of scleral appearance may be guided by various other factors aside from self-domestication, including communicative demands that may relate to irido-scleral contrast as well as the need for efficient photoprotection of the external eye [3, 4, 6, 12]. Due to scaling effects and energetic constraints, larger-bodied mammals expose larger portions of the bulbar conjunctiva during glancing and rely on movements of the eyeball rather than the head to visually scan their surroundings [10]. Therefore, large species with typically bigger eyes [28] might be expected to show stronger pigmentation than small-bodied ones to more effectively protect their ocular epithelia from UV radiation, regardless of their domestication status.

Here, we examined ocular pigmentation in 13 domesticated mammal species compared to their closely related wild relatives to address whether conjunctival depigmentation actually represents a correlate of domestication. Furthermore, we test whether increased conjunctival pigmentation is a correlate of eye size within our species sample. Subsequently, we discuss the implications of our findings for understanding mammalian ocular phenotypes in general, including those of (human) primates.

## Material and Methods

We quantified scleral appearance and irido-scleral contrast in 26 mammalian lineages. These encompassed 13 domesticated groups and 13 representatives of closely related, non-domesticated taxa (Tab. 1). For each of these lineages, we collected at least 15 high-quality photographs that we used for analysis (compare [29]). The photos had to allow for an unambiguous distinction between the iris and the sclera/bulbar conjunctiva. We only included photos showing animals that appeared to be adult, as it has been shown, at least in primates, that scleral pigmentation undergoes ontogenetic changes, with juveniles exhibiting brighter scleral appearance [11]. Our sample is summarized in Table 1, with web links to individual pictures being included in Supplementary Table 1.

**Tab. 1:** Measurements of scleral brightness and highest irido-scleral contrast (HC) in selected domesticated and wild mammals, as well as published data on their species-specific axial ocular diameters. In case ocular diameter measurements were not available, values for similarly sized related species were used (anoa, European rabbit) or the corresponding measurements of the domesticated (polecat, European wildcat, Przwewalski’s horse, wild sheep) or wild (domestic yak) counterpart were adopted. Note that there is an ongoing debate as to whether and to which extent Przewalski’s horses are feral descendants of an ancient domesticated lineage [37]. Data on adult ocular diameter measurements derive from: a - species mean values from [28], b - [38], c - [39], d - [40], e - [41], f - [42]. *value corresponds to *Sylvilagus audobonii*, **value corresponds to *Hippotragus niger*, a bovid of similar body mass, ***value corresponds to *Bos bison*.

| Species |  | N |  | Mean scleral brightness |  | HC |  | Ocular diameter (mm) |  |
| --- | --- | --- | --- | --- | --- | --- | --- | --- | --- |
| Domesticated | Wild | Dom. | Wild | Dom. | Wild | Dom. | Wild | Dom. | Wild |
| Domestic rabbit (*Oryctolagus cuniculus* f. *domestica*) | European rabbit (*Oryctolagus cuniculus*) | 15 | 16 | 204.97 | 184.75 | 184.53 | 168.44 | 18.1<sup>a</sup> | 14.9<sup>b*</sup> |
| Ferret (*Mustela furo*) | Polecat (*Mustela putorius* & *Mustela eversmanni*) | 15 | 15 | 107.9 | 124.6 | 113.07 | 132.67 | 7.5<sup>a</sup> | 7.5<sup>a</sup> |
| Dog (*Canis familiaris*) | Wolf (*Canis lupus*) | 25 | 25 | 171.86 | 142.38 | 122.2 | 84.28 | 20.1<sup>a</sup> | 22.6<sup>a</sup> |
| House cat (*Felis catus*) | European wildcat (*Felis silvestris*) | 16 | 15 | 161.44 | 169.30 | 49.94 | 56.93 | 21.9<sup>a</sup> | 21.9<sup>a</sup> |
| Domestic donkey (*Equus asinus*) | Zebra (*Equus grevyi* & *Equus quagga*) | 21 | 15 | 81.24 | 50.93 | 62.38 | 27.80 | 39.0<sup>b</sup> | 42.5<sup>a</sup> |
| Domestic horse (*Equus ferus caballus*) | Przewalski's horse (*Equus ferus przewalskii*)* | 18 | 15 | 93.06 | 72.67 | 79.06 | 51.93 | 38.7<sup>a</sup> | 38.7<sup>a</sup> |
| Llama (*Lama glama*) | Guanaco (*Lama guanicoe*) | 16 | 15 | 68.41 | 58.27 | 42.56 | 57 | 35<sup>a</sup> | 36<sup>a</sup> |
| Domestic pig (*Sus domesticus*) | Wild boar (*Sus scrofa*) | 16 | 15 | 169.19 | 75.67 | 120.50 | 47.73 | 23.9<sup>c</sup> | 24.8<sup>a</sup> |
| Domestic water buffalo (*Bubalus bubalis*) | Anoa (*Bubalus depressicornis* & *Bubalus quarlesi*) | 15 | 15 | 56.27 | 57.97 | 55.27 | 55.07 | 33.4<sup>d</sup> | 30<sup>a**</sup> |
| Taurine cattle (*Bos taurus*) | European bison (*Bos bonasus*) | 19 | 16 | 91.95 | 99.31 | 93.05 | 111.88 | 30.8<sup>a</sup> | 36.8<sup>e***</sup> |
| Domestic yak (*Bos grunniens*) | American bison (*Bos bison*) | 15 | 20 | 85.07 | 90.13 | 95.2 | 99.35 | 36.8<sup>e***</sup> | 36.8<sup>e</sup> |
| Domestic sheep (*Ovis aries*) | Wild sheep (*Ovis ammon* & *Ovis vignei*) | 17 | 15 | 126.06 | 78 | 82.59 | 53.73 | 26.1<sup>a</sup> | 26.1<sup>a</sup> |
| Domestic goat (*Capra hircus*) | Wild goat (*Capra falconeri*, *Capra ibex*, & *Capra nubiana*) | 18 | 15 | 96.36 | 72.53 | 52.33 | 62.07 | 30.1<sup>f</sup> | 35.0<sup>b</sup> |

We attempted to sample the closest living wild relatives of each domestic species. However, for some lineages this was not possible because too few photos meeting our criteria were available online (Table 1; Supplementary Table 1). This was the case, for instance, for the wild water buffalo (*Bubalus arnee*), wild yak *(Bos mutus*), and African wild ass (*Equus africanus*). In some cases, for the same reason, it was necessary to pool data from more than one closely related wild species to reach our sampling criterion (Table 1; Supplementary Table 1). For example, we sampled both Grevy’s zebras (*Equus grevyi*) and plains zebras (*Equus quagga*) as wild counterparts of the domestic donkey (*Equus asinus*). We did not sample brachycephalic breeds of domesticated mammals, since they are known to frequently exhibit numerous ocular pathologies that can result in aberrant pigmentation (see e.g., [30]). We also excluded albinotic individuals and did so for partially leucistic/piebald individuals if the integument surrounding the eye was affected by depigmentation.

To quantify ocular pigmentation, we extracted greyscale luminance values from the pictures using the plot profile function in ImageJ [31], as described in [6] and [11]. For both the visible scleral and the iridal portions of the eye, we noted the highest and lowest greyscale values found in a given individual. We then used these values to calculate the highest contrast (HC) between these tissues and to approximate the mean brightness of the scleral portion of the visible eye (= average between lowest and highest measured grey value luminance in the sclera). Reflections, shadows, and the corneal limbus, the pigmentation of which might form a strong contrast with the adjacent tissues [3], were carefully avoided. For each picture, only one eye was sampled even when both eyes of an individual were visible. In such cases, we chose the better illuminated eye.

Statistics were created in R [32] and performed on the means calculated for each species. Pagel’s *λ* was used to measure the phylogenetic signal in the data. We applied phylogenetic paired *t*-tests (*phyl.pairedttest* function in the phytools package; [33]) to test for an effect of domestication on log-transformed scleral brightness and iridoscleral contrast values. To determine the potential influence of ocular diameter on scleral brightness, we applied phylogenetic generalized least squares (PGLS) regression (form: log(scleral brightness)~log(ocular diameter):domestication status, correlation structure: Pagel’s *λ*; model fit: maximum likelihood). Data on axial ocular diameter were retrieved from the literature (see legend for Tab. 1 for details on sources). Normal distribution of data as well as of model residuals was checked using the Shapiro-Wilk test. Phylogenetic tree topology and species-level divergence dates were derived from VertLife.org [34], with additional divergences between domesticated and wild forms dated according to [35] and [36] if not included in the VertLife database.

## Results

We did not find marked signatures of domestication in the conjunctival pigmentation of the species pairs studied (Fig. 1). Scleral brightness was generally high in lagomorphs and carnivorans and low in ungulates, regardless of whether the population in question was domesticated or not. Accordingly, scleral brightness in our overall dataset exhibited a notable phylogenetic signal (Pagel’s *λ* = 0.75, likelihood ratio test *p* < 0.001), and the same was true for iridoscleral contrast (Pagel’s *λ* = 0.80, likelihood ratio test *p* < 0.001). Phylogenetic paired t-tests revealed that both scleral brightness (*t* = 0.953, *p* = 0.363) and irido-scleral contrast (*t* = 0.956, *p* = 0.362) are not significantly different between domesticated lineages and their wild counterparts. Thus, domesticated mammals generally resemble their wild ancestors in terms of scleral brightness and do not share a uniform phenotype of conjunctival pigmentation (Fig. 2A). On average, scleral brightness was only moderately higher in domesticated (mean scleral brightness: 116.4; SD: 44.4) compared with wild lineages (mean scleral brightness: 98.2; SD: 42.0; Fig. 2B), although the differences were pronounced in some species pairs, such as in pigs. Here, domestic pigs (*Sus domesticus;* mean scleral brightness = 169.19) tended to show highly depigmented conjunctivae, whereas those of wild boar were typically dark (*Sus scrofa;* mean scleral brightness = 75.67; Fig. 1).

**Fig. 1:**
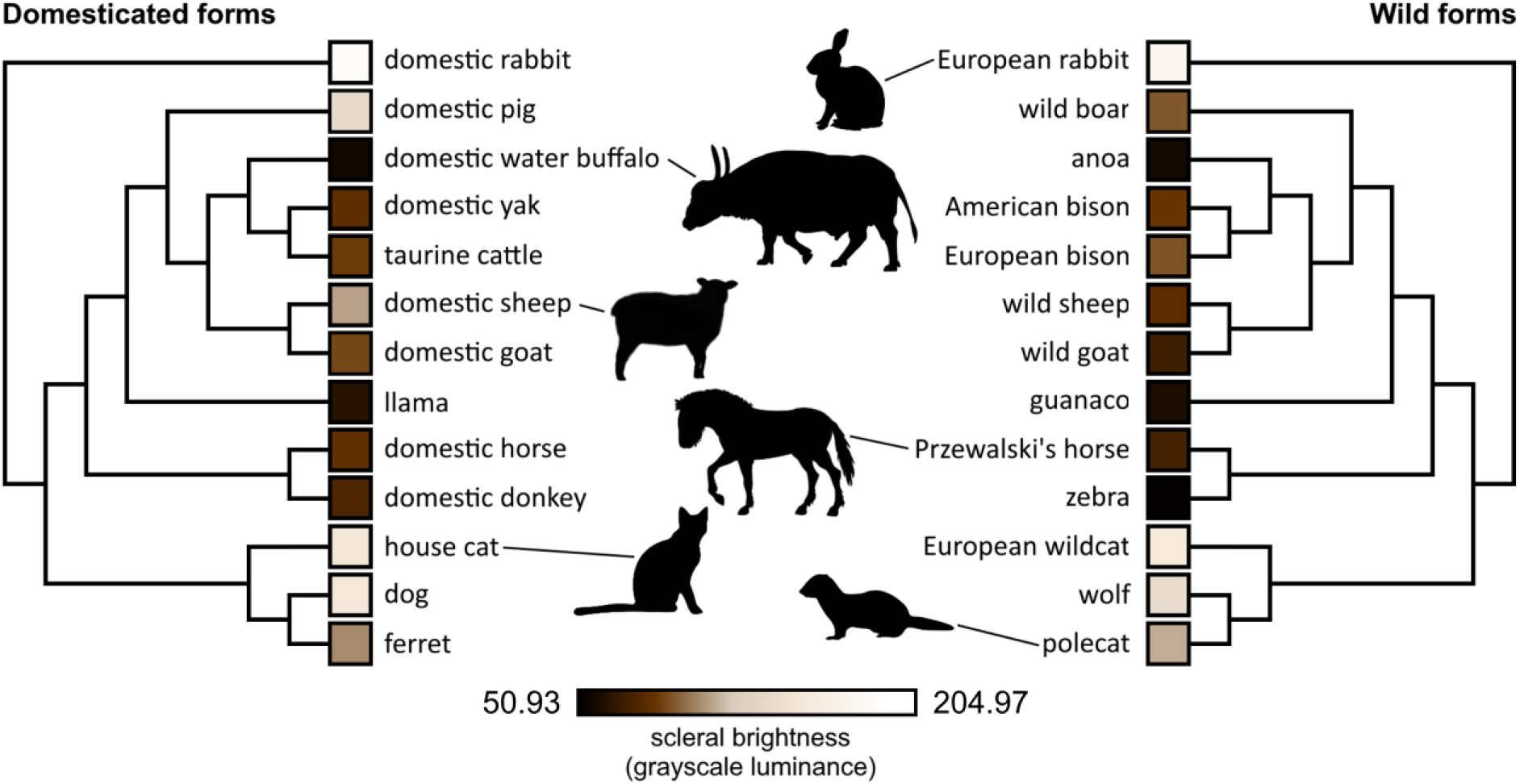
Phylogenies with annotated scleral brightness values (grayscale luminance, colors only approximated) for the domesticated species (left) and the respective wild forms (right) featured in this study. Silhouette credits: Przewalski’s horse by Mercedes Yrayzoz, domestic sheep by Gabriela Palomo-Munoz, European rabbit by Anthony Caravaggi, domestic water buffalo by Cristopher Silva, others in public domain. All silhouettes derive from PhyloPic.

**Fig. 2:**
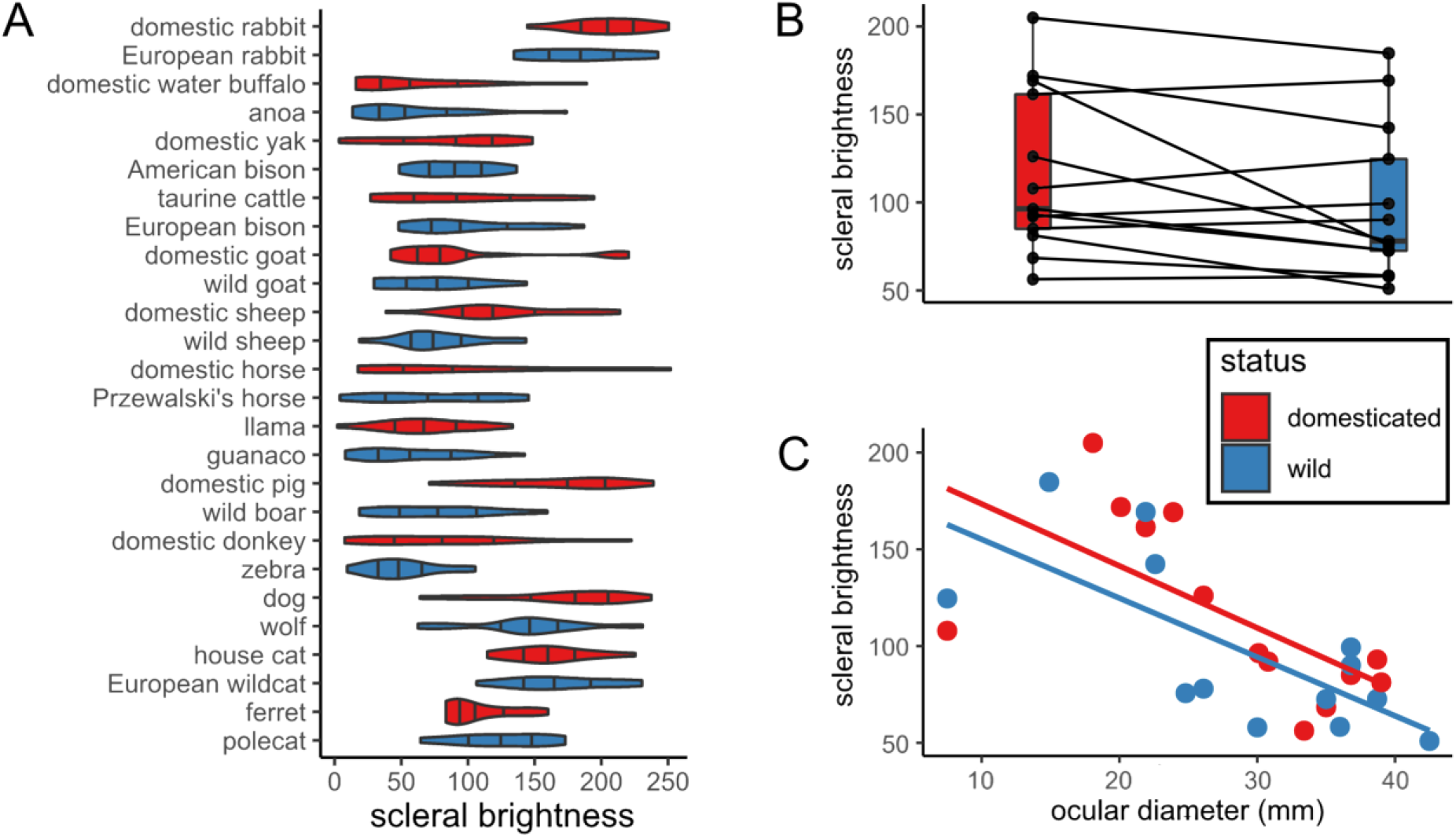
Scleral brightness (approximated by grey values of the scleral portion of the externally visible eye and thus including the conjunctiva) in closely related wild and domesticated mammals. A: Distribution of scleral brightness values in the studied species. Note that domesticated and wild forms of each species mostly resemble each other. B: Paired boxplot showing scleral brightness in domesticated forms compared to their close wild relatives. C: Plot of scleral brightness as a function of ocular diameter. Note that species with larger eyes tend to exhibit greater degrees of conjunctival pigmentation and thus lower scleral brightness.

We found no significant effects of eye size on scleral brightness in an initial PGLS model that included the entire species sample (*p* > 0.38, *n*_taxa_ = 26, Table 2). However, when inspecting the data, we noticed that the species pair with the smallest eyes, ferrets and polecats, represented marked outliers (Fig. 2C). When those were removed from the sample, a significant negative effect of eye size on scleral brightness emerged for both domesticated and wild forms. In this reduced sample, larger-eyed species displayed significantly greater conjunctival pigmentation (*p* < 0.05, *n*_taxa_ = 24, Table 2).

**Tab. 2:**
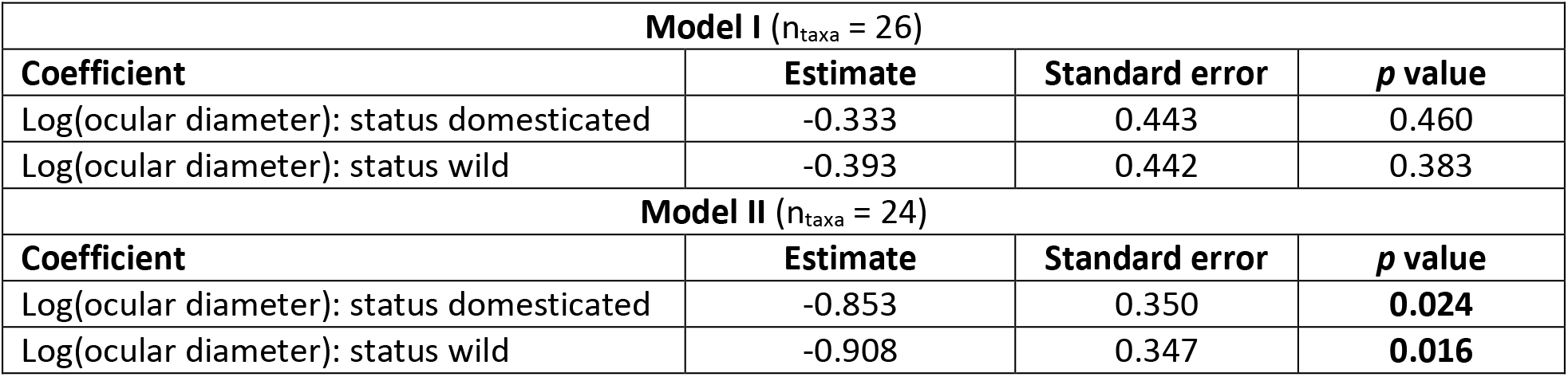
Comparison of PGLS statistics (scleral brightness in relation to eye size and domestication status) including (Model I) and excluding (Model II) the genus *Mustela* (ferrets and polecats).

| **Model I ($n_{\text{taxa}} = 26$)** |  |  |  |
| --- | --- | --- | --- |
| **Coefficient** | **Estimate** | **Standard error** | **p value** |
| Log(ocular diameter): status domesticated | -0.333 | 0.443 | 0.460 |
| Log(ocular diameter): status wild | -0.393 | 0.442 | 0.383 |
| **Model II ($n_{\text{taxa}} = 24$)** |  |  |  |
| **Coefficient** | **Estimate** | **Standard error** | **p value** |
| Log(ocular diameter): status domesticated | -0.853 | 0.350 | **0.024** |
| Log(ocular diameter): status wild | -0.908 | 0.347 | **0.016** |

## Discussion

Our results suggest that scleral appearance in most domesticated mammals is not notably different from that of closely related wild species. Instead, this trait seems to be largely unaffected by the domestication process and thus should not be considered part of the mammalian domestication syndrome (contra [16]). In this regard, conjunctival depigmentation differs from both integumental and iridal pigmentation [2, 18]. Different evolutionary drivers might thus determine the expression of these traits, which are probably affected by the different developmental trajectories of melanocyte populations in the skin, uvea, and conjunctiva [25, 26, 43]. Interestingly, in taxa where scleral appearance is rather static, ocular appearance and facial expressions mediated by the eyes can nevertheless change radically through the domestication process, as exemplified by dogs in comparison to wolves [44]. In some domesticated mammals, the degree of conjunctival pigmentation varies greatly between breeds. Appaloosa horses, for instance, are known to have depigmented conjunctivae [45], deviating from the dark eyes of most other horse breeds. This variation bears an important caveat for our study: since we could not rigorously control for breed representation in our photo sample, certain breed-specific phenotypes may be over- or underrepresented. This phenotypic variability, although a source for biases in studies such as ours, may prove to be greatly valuable for future research on the evolution of ocular pigmentation in mammals. Genomic comparisons between dark and light-eyed breeds could help determine the genetic basis of these traits and thus help us understand the evolution of human ocular appearance. While the genes controlling coat coloration in domesticated animals have been extensively studied [23], the genetic determinants of scleral appearance have so far remained completely unexplored.

Although we did not recover statistically significant differences between the scleral appearance of domesticated and wild forms at the level of our full sample, some domesticated lineages such as pigs and sheep indeed show conspicuously brighter eyes than their wild relatives. At the moment, we cannot exclude that this is a pleiotropic byproduct of domestication in these specific groups. Potential adaptive functions of reduced conjunctival pigmentation in these species are not apparent. Interestingly, a brighter scleral appearance seems to be generally typical for juvenile mammals (e.g., [8, 11]). Therefore, the brighter eyes could align with several other traits deemed indicative of paedomorphosis in some domesticated lineages [36]. But do we need to invoke impaired migration of embryonic neural crest cells to explain this pigmentation pattern?

At the cellular level, depigmentation of the conjunctiva can be achieved in two ways. First, the density of melanocytes could be reduced. This would be consistent with the general predictions of the domestication syndrome hypothesis. Second, conjunctival melanocytes could still be abundant in the tissue but no longer produce enough melanin to impose a macroscopic effect. Of course, a combination of these two factors is also conceivable. It is important to note that in at least some mammals, even fully transparent conjunctivae contain melanocytes. This contrasts with the absence of these cells in the depigmented skin areas of domesticated and alleged self-domesticated species with pied coats [19]. Apart from humans, the presence of pigment-bearing cells in macroscopically transparent conjunctivae has been demonstrated in marmosets and capuchins [15]. In response to as yet unknown stimuli, these cells may overproduce pigment, resulting in brownish patches on the conjunctival epithelium (intraepithelial nonproliferative melanocytic (hyper)pigmentation, not to be confused with melanoma – Jakobiec [46]). Thus, one cannot simply equate a decrease in pigmentation with a quantitative reduction of melanocytes in the conjunctiva. To test the different evolutionary scenarios, the abundance of melanocytes in species differing in conjunctival pigmentation and/or domestication status needs to be mapped in a comparative fashion. Unfortunately, this has not yet been accomplished.

Ecological factors have only recently gained attention in discussions on the evolution of ocular pigmentation in mammals, with arguments being made for an important role of conjunctival pigmentation in photoprotection [3, 12]. Corneal stem cells located at the limbus are particularly vulnerable to UV radiation and are likely to benefit from melanin shielding [3], as is the conjunctival epithelium itself. Data on primates suggest that the degree of habitual eye ball rotation, and thus radiation exposure of the conjunctiva, correlates positively with body size ([10]; and thereby also eye size - [38]). If this pattern is applicable to mammals in general (as anecdotal observations might suggest, compare [47]), it would fit well with our preliminary finding that darker conjunctivae are characteristic of large-eyed species. Interestingly, ferrets and polecats, animals with barely exposed scleral portions of the eyeball and the smallest eyes within our sample, do not comply to the above scheme. They had to be excluded from the analysis to yield a significant correlation between eye size and conjunctival pigmentation. This could simply be related to scaling effects on scleral tissue thickness and thus translucency (note that the scleral overlays the well vascularized uvea and strongly pigmented retina), but comparative data are needed before any reasonable conclusions can be drawn. In any case, our sample is obviously too small and too narrow in terms of phylogenetic and ecological representativeness to derive general patterns for mammals. An expanded dataset is required to robustly test whether eye dimensions are positively correlated with conjunctival pigmentation and to robustly infer its potential adaptive significance. We would also like to point out that eye movements in mammals with forward-facing eyes, for instance primates and cats, are fundamentally different from those found in groups such as ungulates and lagomorphs [48]. For more specific comparative studies, such morphological differences need to be considered along with species-specific activity rhythms.

What are the implications of our findings for understanding the evolution of ocular pigmentation in primates, including humans? First and foremost, they challenge the notion that species such as marmosets, bonobos, and humans acquired their depigmented conjunctivae through self-domestication [7, 11, 16], even if they should have experienced such a process in their evolutionary history. Instead, secondarily depigmented eyes may have evolved to facilitate communication mediated by eye-gaze ([4, 14] but see [6] for a rebuttal of this idea) or other forms of social signaling. Specifically, in humans, depigmented conjunctivae could represent a sexually selected trait, and potential effects of genetic drift on scleral appearance still need to be appraised [6]. It should also be pointed out that depigmented conjunctivae in different primate taxa may have emerged due to different evolutionary pressures (or a lack thereof), given the in parts great phylogenetic and morphological disparities between them. Our preliminary data on conjunctival pigmentation as a correlate of eye size also raise additional questions about the evolution of the human ocular phenotype: While the transparent conjunctivae of marmosets [7, 12] resemble the ocular phenotype found among other small mammals, such as lagomorphs, the eyes of humans are in striking contrast to those of mammalian species of comparable body and eye size such as ungulates and large-bodied carnivorans [47]. If we assume that conjunctival pigmentation does indeed adaptively shield exposed ocular epithelia from UV radiation, how do humans (and some other great apes such as Sumatran orangutans - [6, 12]), which are strictly diurnal animals and evolved at low latitudes, compensate for its reduction? So far, this interesting question has attracted little scientific attention (but see [3] for relevant discussions on the distribution of limbal stem cells in the human eye).

Comprehensive comparative ophthalmological studies are needed to place human ocular pigmentation into its phylogenetic context and to identify its evolutionary drivers. Due to their availability to researchers, domesticated mammals may importantly contribute to our understanding of these factors in the future. The domestication process itself, however, seems to not have markedly shaped scleral appearance in mammals, suggesting that the self-domestication hypothesis is not the key to understanding the unique traits of the human eye.

